# Spectrograms of kent sounds from seven Ivory-billed Woodpecker expeditions show 587 Hertz as a pattern

**DOI:** 10.1101/2024.05.07.592969

**Authors:** John D Williams

## Abstract

Although the Ivory-billed Woodpecker is widely considered to be extinct, expeditions led by professional ornithologists have acquired visual, video, and audio evidence since 1968, with increases in data since 2000. Audio evidence for “kent” sounds (n=150) were examined for commonality; this was found to be true with a pattern at 587 Hz that suggests a morphological basis. Differences from other species were found on spectrograms. The patterns extend to different states and years and reinforces that the species persists.

## Introduction

The Ivory-billed Woodpecker (Campephilus principalis) (IBWO) has been observed sporadically for over two hundred years. Observations by prominent scientists, explorers, naturalists, hunters and other individuals differ and are sometimes contradictory. There is much that is not known about this species, its behavior, and ecological needs. It is not well documented in photographic images, videos, or audio recordings. Although there is a body of evidence– treated forensically and with morphometric analysis and peer review– since 2000 from professional and significant search efforts, the results are not considered conclusive enough for a layperson to consider the species to be extant [1]. In popular literature for this, the “definitive image” is highly sought.

However, birds and other animals can be identified by sound. The IBWO has been observed to produce a variety of sounds including knocking on trees as a signal (a “double knock”), calls produced at the nest that are thought to be from stressed birds [2], and conversation calls known as “kents”. In a phonetic sense, kents have been written in the literature as kent, kint, kaint, haint, hant, heent, keent and other interpretations [3]. This most likely reflects a wider range of sounds for this species than is commonly accepted. All of these sounds have a nasal quality.

Because the kent sound can also be used as a descriptor for other species, it is necessary to use quantitative study from spectrograms to look for differences. Spectrograms are visual representations of sounds, relating time to Hertz frequency. For example, a kent sound thought to be made by an Ivory-billed Woodpecker, as seen on a spectrogram is shown here. These kent sounds are characterized by horizontal lines in a ladder-like structure. Virtually all of the kent sounds thought to be from the IBWO that are examined in this paper look like this:

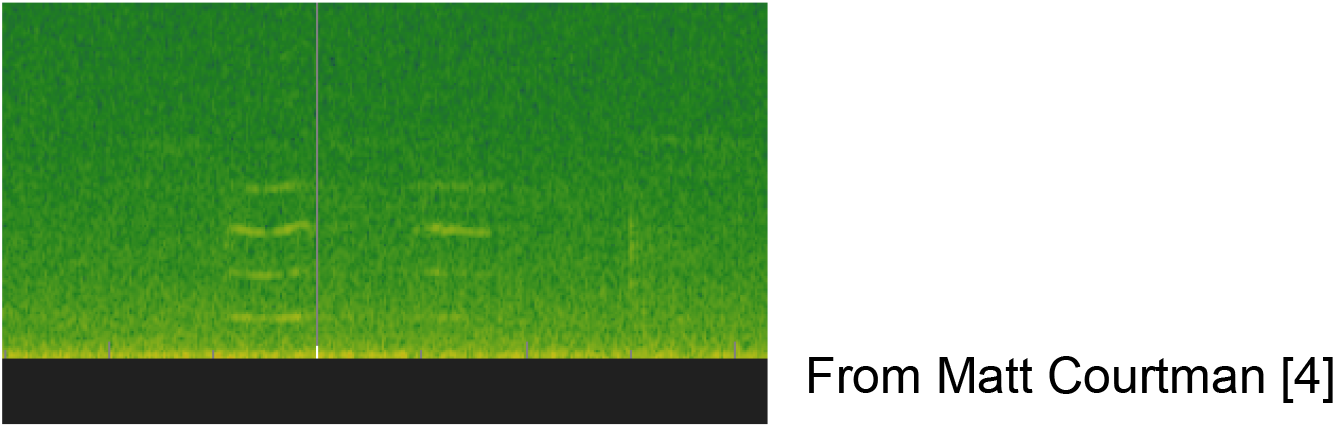

From Matt Courtman [4]

These kent sounds also compare well– flattish lines, spacing between lines, duration, nasal quality to the ear– with the only visually-correlated recordings for this species, obtained by Arthur Allen and Paul Kellogg of Cornell, in 1935, an example here [5]:

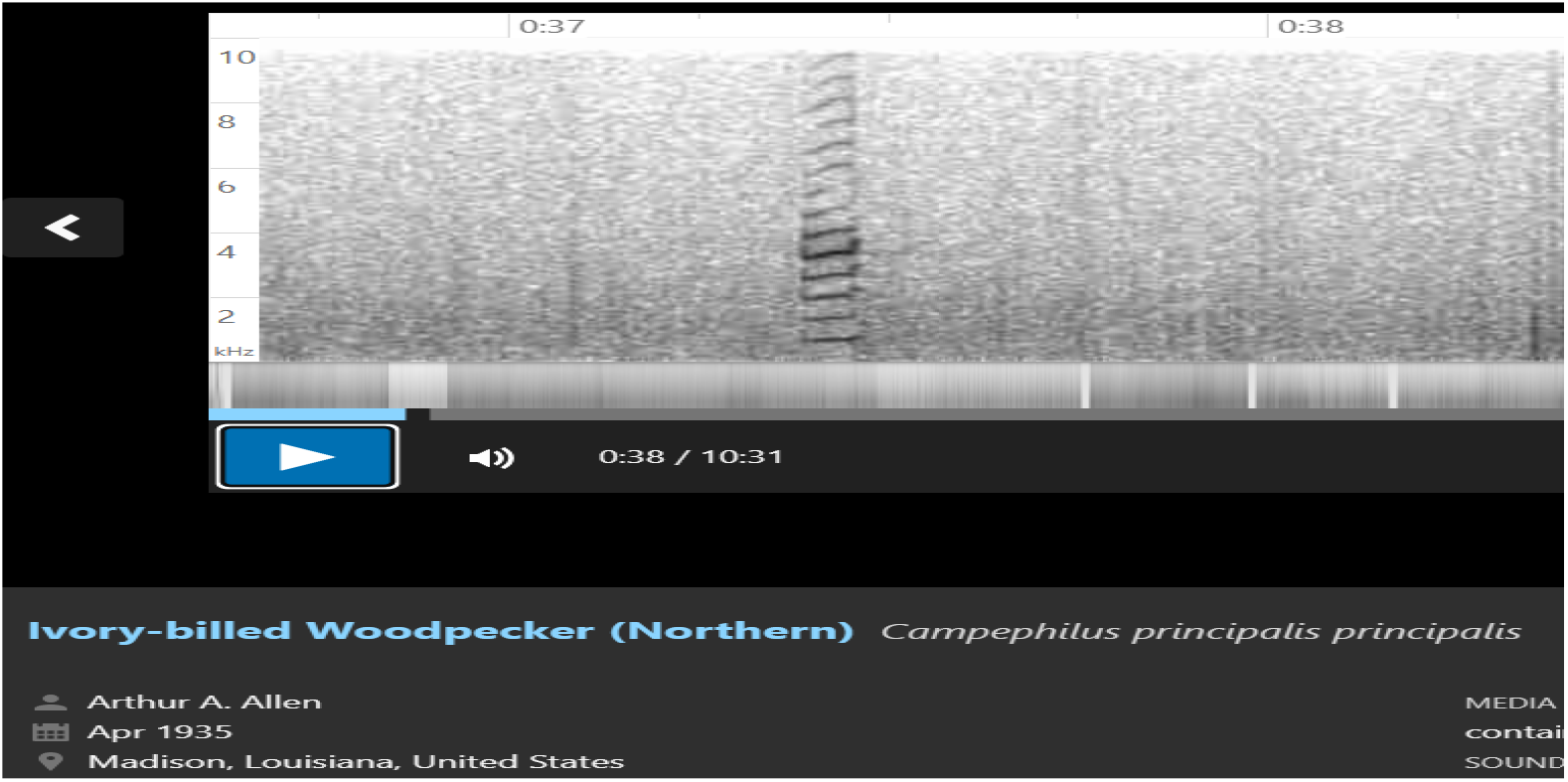

However, the Allen-Kellogg sounds, if described phonetically, are not “kents”, but described to be from birds stressed by human presence at an active nest, and thus making different sounds which have more complexity [2]. They are not part of the dataset used for this paper.

Qualities that kents have, and comparable spectrograms, are discussed in a Project Principalis--National Aviary writing, in the section Vocalizations [6]. https://www.aviary.org/conservation/project-principalis/natural-history/sounds/

## Materials and Methods

For all kent sounds in this study, the website program Sonic Visualizer [7] was used to create and examine spectrograms. This program was chosen for ease of use, clarity, precision, and comparison with other IBWO research. Spectrogram analysis uses time as the X axis and vibrations per second (Hertz or Hz) as the Y axis, as seen here on Sonic Visualizer:

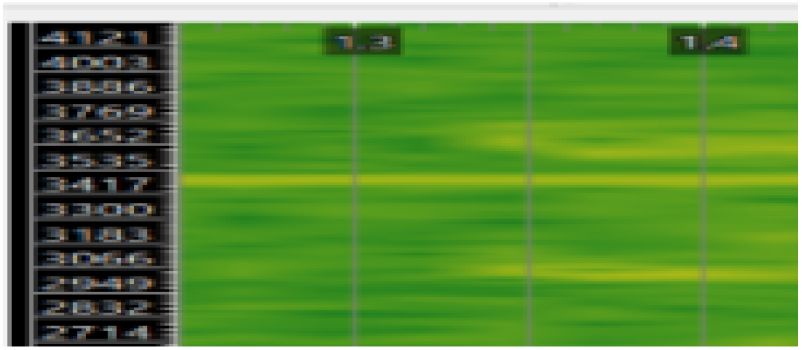

Original sound recordings were not altered. To standardize, adjustments were made to the x and y axes to see length of calls and known or reported Hz frequency ranges. Comparisons were made to existing data for other members of the same genus Campephilus by using the sound libraries Macaulay Library [5] and Xeno-canto [8].

The program Online Tone Generator [9] was used for matching kent sounds to known Hz, which is comparable to listening to a note on a piano and having it identified numerically. Estimated accuracy for this is +/- 10 Hz.

To clearly show relationships of harmonic and non-harmonic spectrogram Hz frequencies, visual data was changed to number tables directly by this method:

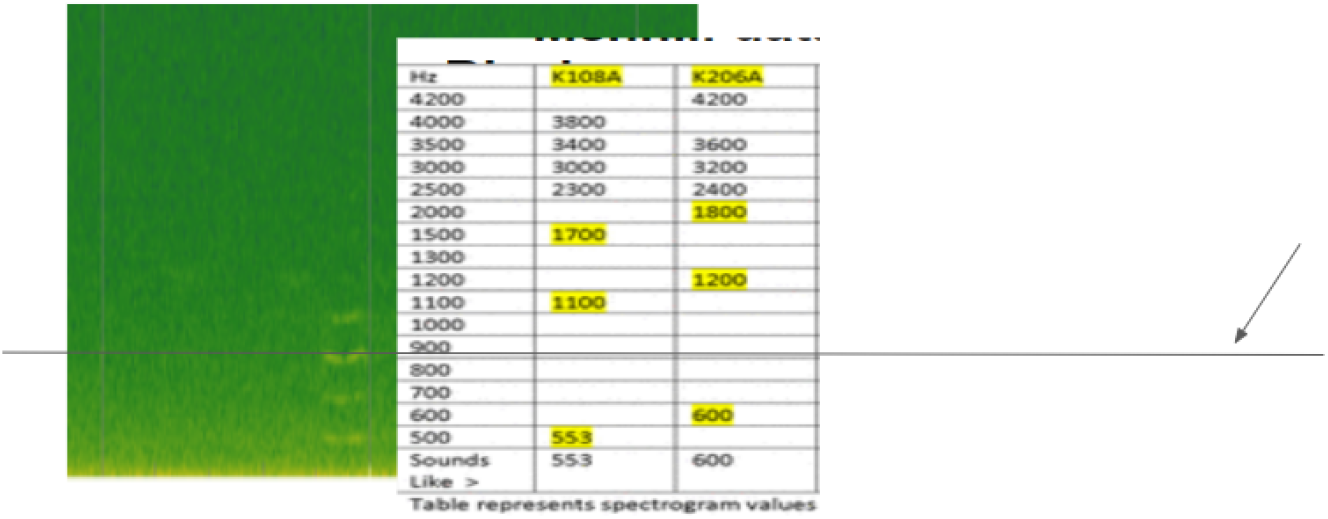

Method of changing spectrogram directly to number table. Arrowed line represents 900 Hz

Using these methods, approximately 150 different kent sounds were compared. All the sounds were obtained from expeditions, involving professionals in various aspects of field biology, focused on locating evidence for the Ivory-billed Woodpecker. Recordings were obtained either from handheld devices or autonomous recordings units (ARUs).

## Results and Discussion

### 1. Quantitative differences between IBWO kents and other kentlike sounds

In order for sounds to be considered as kents from the Ivory-billed Woodpecker, it is necessary to distinguish them from other species that make similar sounds. An often-repeated critique is that it is not easy to tell IBWO kents from similar sounds made by nuthatches (genus <u>Sitta</u>), Bluejay (<u>Cyanocitta cristata</u>), or a small number of other species. However, sounds suspected to be from the IBWO show important spectrogram traits that can differentiate them from these and other species.

Kentlike sounds are converted from spectrograms and compared below using data from the Auburn University-University of Windsor dataset [10]. This was obtained with Geoff Hill’s Choctawatchee River expedition in December 2005 [11]. Nuthatch and Bluejay kentlike sounds were obtained from the best matches found in Xeno-canto [8]. It was found that “octave harmonics”, multiples of a Hertz frequency, were common in sounds suspected to be from the IBWO.

| Hz<br>v | K108A | K206A | K206B | Ke1353 | Ke1018 | WBNH | RBNH | BJ | BJ |
| --- | --- | --- | --- | --- | --- | --- | --- | --- | --- |
| 4200 |  | 4200 |  |  |  |  |  |  |  |
| 4000 | 3800 |  |  |  |  |  |  |  |  |
| 3500 | 3400 | 3600 |  |  |  |  | 3500 |  |  |
| 3000 | 3000 | 3200 |  | 3100 | 3200 | 3200 | 3100 | 3200 |  |
| 2500 | 2300 | 2400 | 2400 | 2700 | 2500 | 2800 | 2500 |  | 2500 |
| 2000 |  | 1800 | 1800 | 2200 |  | 2000 | 2200 | 2300 |  |
| 1500 | 1700 |  |  | 1500 | 1600 |  | 1600 | 1600 | 1600 |
| 1300 |  |  |  |  |  | 1400 |  |  |  |
| 1200 |  | 1200 | 1200 |  |  |  |  |  |  |
| 1100 | 1100 |  |  | 1100 |  |  | 1100 |  |  |
| 1000 |  |  |  |  |  |  |  |  |  |
| 900 |  |  |  |  |  |  |  |  |  |
| 800 |  |  |  |  | 836 |  |  |  | 845 |
| 700 |  |  |  |  |  |  |  | 794 |  |
| 600 |  | 600 | 613 |  |  |  |  |  |  |
| 500 | 553 |  |  | 546 |  |  |  |  |  |
| Sounds like | 553 | 600 | 613 | 546 | 836 | 1400 | 1100 | 794 | 845 |

Table represents spectrogram values in Hz. IBWO info used with permission from Dan Mennill [10]. K and Ke represent putative IBWO kents. WBNH= White-breasted Nuthatch RBNH= Red-breasted Nuthatch BJ= Bluejay

Octave harmonics in yellow. WBNH and RBNH have Hz spectrogram line between first octave (red).

Additional kentlike sounds from the Auburn-Windsor audio recordings were examined. All of these sounds are suspected to be from the Ivory-billed Woodpecker. Again the pattern of octave harmonics is dominant.

| Hz v | Kent 3-13<br>1046 | Kent 3-15<br>1007 (1) | Kent 3-15<br>1007 (2) | Kent 3-15<br>1007 (3) | Kent 3-15<br>1007 (4) | Kent 3-16<br>1626 | Kent 3-18<br>1124 (1) | Kent 3-18<br>1124 (2) | Kent 3-18<br>1124 (3) |
| --- | --- | --- | --- | --- | --- | --- | --- | --- | --- |
| 4200 |  |  |  |  |  |  |  |  |  |
| 4000 |  |  | 4250 | 4250 |  |  |  |  |  |
| 3500 | 3596 |  |  |  |  |  |  |  |  |
| 3000 |  |  | 3400 | 3400 |  |  |  |  |  |
| 2500 | 2850 |  | 2500 | 2500 | 2900 | 2550 |  | 2600 | 2600 |
| 2000 | 2200 | 2400 |  |  | 2200 |  | 2100 |  |  |
| 1500 |  |  | 1600 | 1600 |  | 1900 |  | 1950 | 1700 |
| 1400 |  |  |  |  | 1400 |  | 1400 |  |  |
| 1300 |  |  |  |  |  | 1300 |  | 1300 |  |
| 1200 |  | 1200 |  |  |  |  |  |  | 1200 |
| 1100 |  |  |  |  |  |  |  |  |  |
| 1000 |  |  |  |  |  |  |  |  |  |
| 900 |  |  |  |  |  |  |  |  |  |
| 800 |  |  | 838 | 838 |  |  |  |  | 882 |
| 700 | 700 |  |  |  | 700 |  | 730 |  |  |
| 600 |  |  |  |  |  | missing |  | 660 |  |
| 500 |  | 550faint |  |  |  |  |  |  |  |
| 400 |  |  |  |  |  |  |  |  |  |
| Sounds like > | 741 | 660 | 838 | 838 | 736 | 660 | 730 | 660 | 851 |

Table represents spectrogram values in Hz. Info used with permission from Dan Mennill. Octave harmonics in yellow.

In 1968, the ornithologist John Dennis recorded sounds attributed to the Ivory-billed Woodpecker [12]. Examining these kents with the same process adds the data below. There are a total of 18 kents with these same values. The 18 kents show variation in amplitude for the spectrogram lines but are consistent with Hz frequencies. This again shows octave harmonic structure:

|  |
| --- |
| Hz |
| 2650 to 3000 |
| 2300 |
| 2000 |
| 1700 |
| 1200 |
| 587 Hz (sounds like) |

John Dennis kent recordings (the values occur 18 times in the recording) [12]. Table represents spectrogram values in Hz. Octave harmonics in yellow.

With this precision, sounds suspected to be from the IBWO show important spectrogram traits that can differentiate them from three other species thought to be the source of some reported kents. IBWO kent sounds are harmonic and in octaves.

For example, if an audible sound is at 472 Hz, the lowest spectrogram line would show at 472, and the next line higher will show at close to twice this– an octave harmonic--at around 950 Hz. A third line will be present, an octave above, at 950 + 472, or near 1400 Hz. Michael Collins originally noted this doubling in his spectrogram analysis of ‘high-pitched” calls obtained during observations of an Ivory-billed Woodpecker being harassed by crows in 2006 [13]. Mark Michaels of Project Principalis has recently shown that in related recordings, the third harmonic line is prominent (although not always the Hz frequency heard by a human observer) [14].

Nuthatches can make kentlike sounds. However, on spectrograms, nuthatch species show lines that are non-harmonic. For human singers, this is the equivalent of dissonance, or “bad singing”. This has implications suggesting that for nuthatches, “kents” serve as alarm calls because dissonant sounds are commonly used in Kingdom Animalia for this purpose [15].

The Bluejay can have octave harmonics, but does not produce kentlike sounds with an audible frequency below approximately 800 Hz; this holds true with numerous samples as found on Xeno-canto [8].

Great Blue Herons, Coots, grackles, and some few other species are said to kent. However, recordings of these do not exist on Xeno-canto such that a spectrogram, plus the sound, match at all with putative IBWO data from dedicated expeditions [8].

From these data sources, octave harmonics and a fundamental (lowest line) often below 800 Hz dominate sounds that are thought to be from the Ivory-billed Woodpecker. Recordings of these sounds occurred at times when numerous competent ornithologists and birders reported sightings, in the same area, of this species [11].

Neither nuthatches nor the Bluejay have kentlike calls that match quantitatively, at the spectrogram level, with kent calls recorded by focused Ivory-billed Woodpecker expeditions. This directly addresses the critique of these species being confusing for identification.

This pattern of kentlike sounds having octave harmonics also holds for other species of Campephilus, notably the Magellanic Woodpecker (<u>C. magellenicus</u>) as can be found on Xeno-canto [8].

### 2. Relationships between suspected IBWO sounds from different expeditions

This section compares kent sounds recorded from seven professional efforts directed at finding the Ivory-billed Woodpecker. All the kent sounds used are octave-harmonic. None were reported to be from stressed birds, and none were from birds that were observed during the kent calls. The recordings are from four different states, six different locations, and span 56 years. These are–

1. John Dennis, 1968 Texas [12]
2. Cornell Lab of Ornithology, 2006-2010 Arkansas [16]
3. Auburn University- [University of Windsor, 2005-2006 Florida [10]
4. Project Coyote-Project Principalis with National Aviary, 2007-ongoing Louisiana [6]
5. Matt Courtman, Mission Ivorybill, 2017-ongoing Louisiana [4]
6. Peter Janow, LI Nature Corps with Keith Collins, 2024 Florida [17]
7. Peter Janow, LI Nature Corps with Mission Ivorybill, 2024 Louisiana[18]

Recorded kent sounds were examined for patterns specifically for audible Hertz frequencies. Total kents to work from were 150 with the majority from the Auburn-Windsor dataset, but with considerable numbers from other sources. All were from experienced teams and observers, in many cases professional ornithologists. All expeditions either followed or reported visual sightings. Three obtained visual recordings of differing quality, but able to show fieldmarks, behavior, and quantitative analysis consistent with the IBWO. It is thus valid (and a tribute to the difficult fieldwork involved) to examine these. All recordings were obtained from digital sources; there is no reason nor reference to believe these were altered for speed such that original Hz values were compromised greatly. Both the Auburn-Windsor Florida and Cornell Arkansas recordings were examined previously in their respective bioacoustic university labs.

All sounds used fit the standard of octave harmonics with no dissonant lines as in Part 1 of this Results and Discussion.

My examination of the data began with creating a rough histogram of kent sounds from two groups. It was found that the individual kents formed clusters around certain Hertz frequencies. Each dot represents a single recorded kent sound–

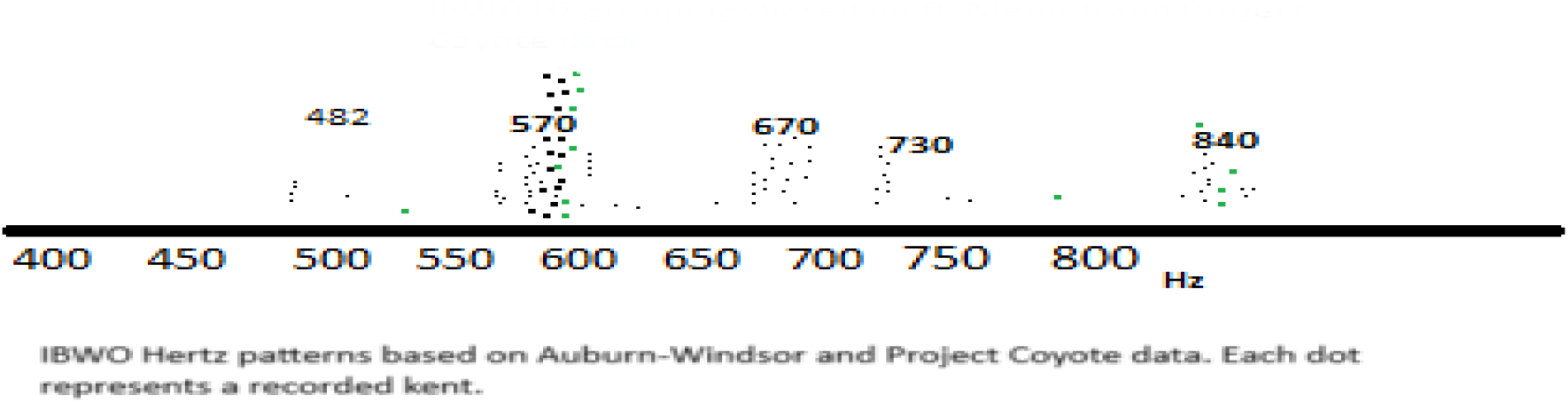

Dots were then added from data obtained by John Dennis and Matt Courtman. Patterns of certain Hertz frequencies were reinforced–

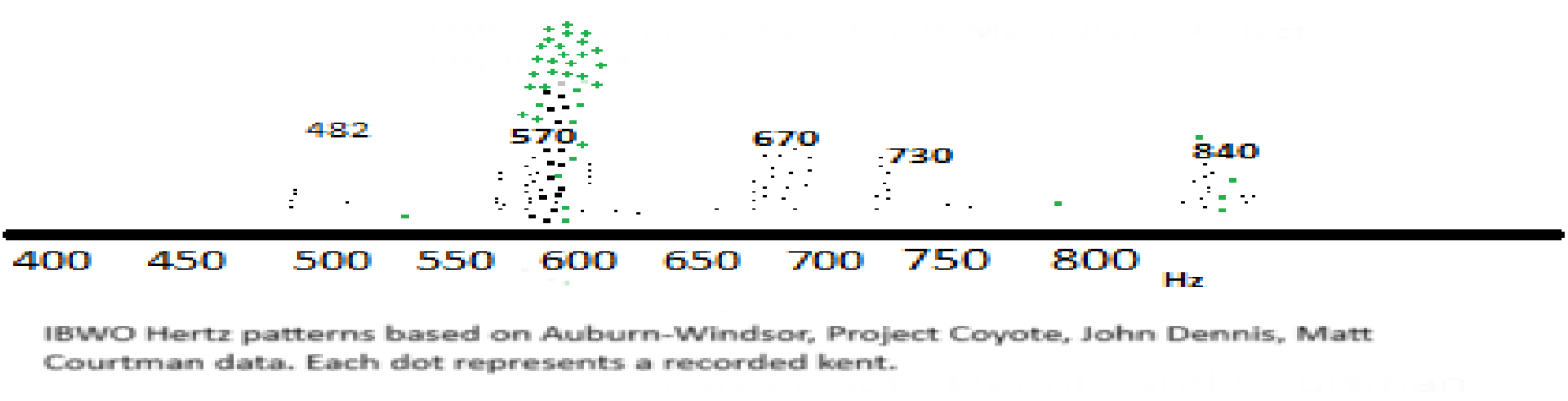

Data points were added from the Cornell efforts in Arkansas–

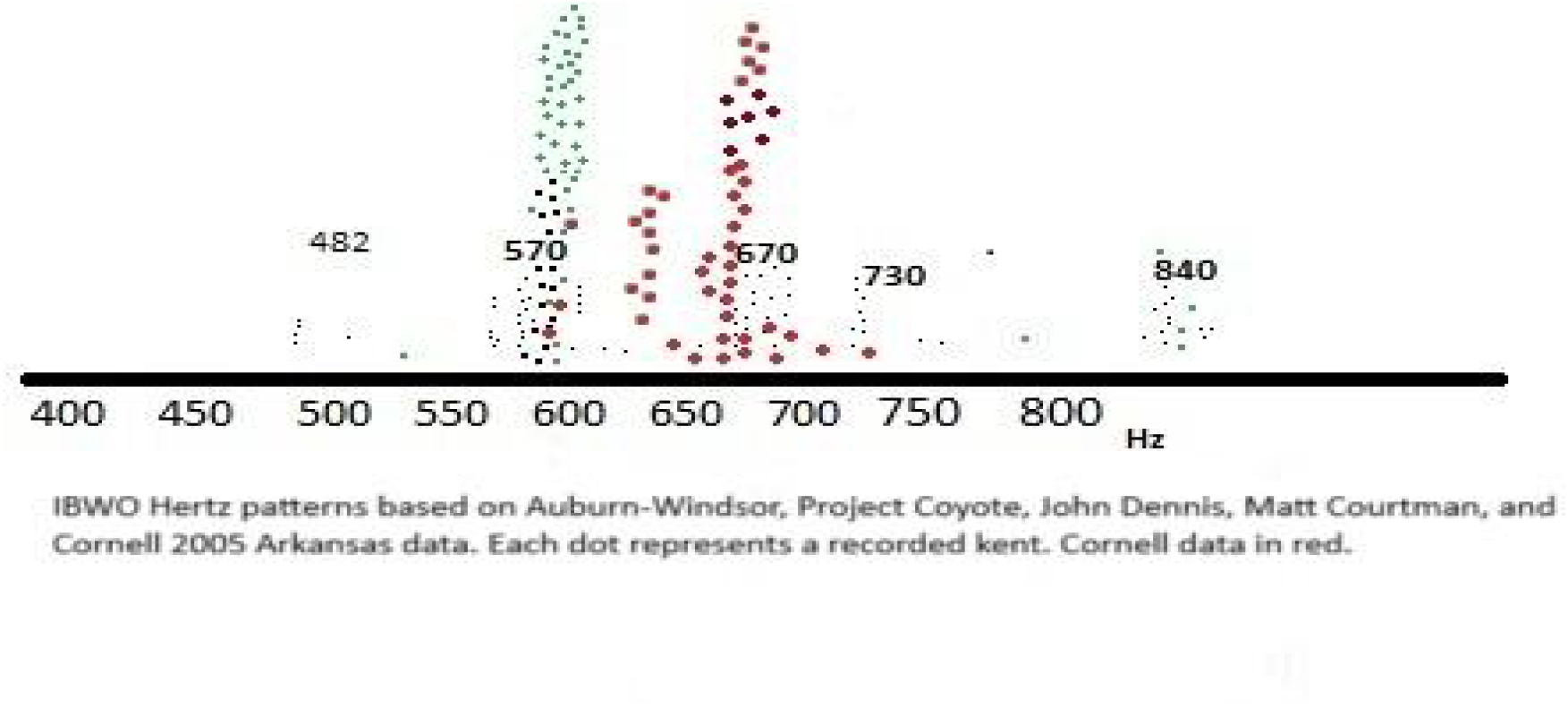

Newer data points obtained in 2024 were added. Finally, a histogram was created from all the data–

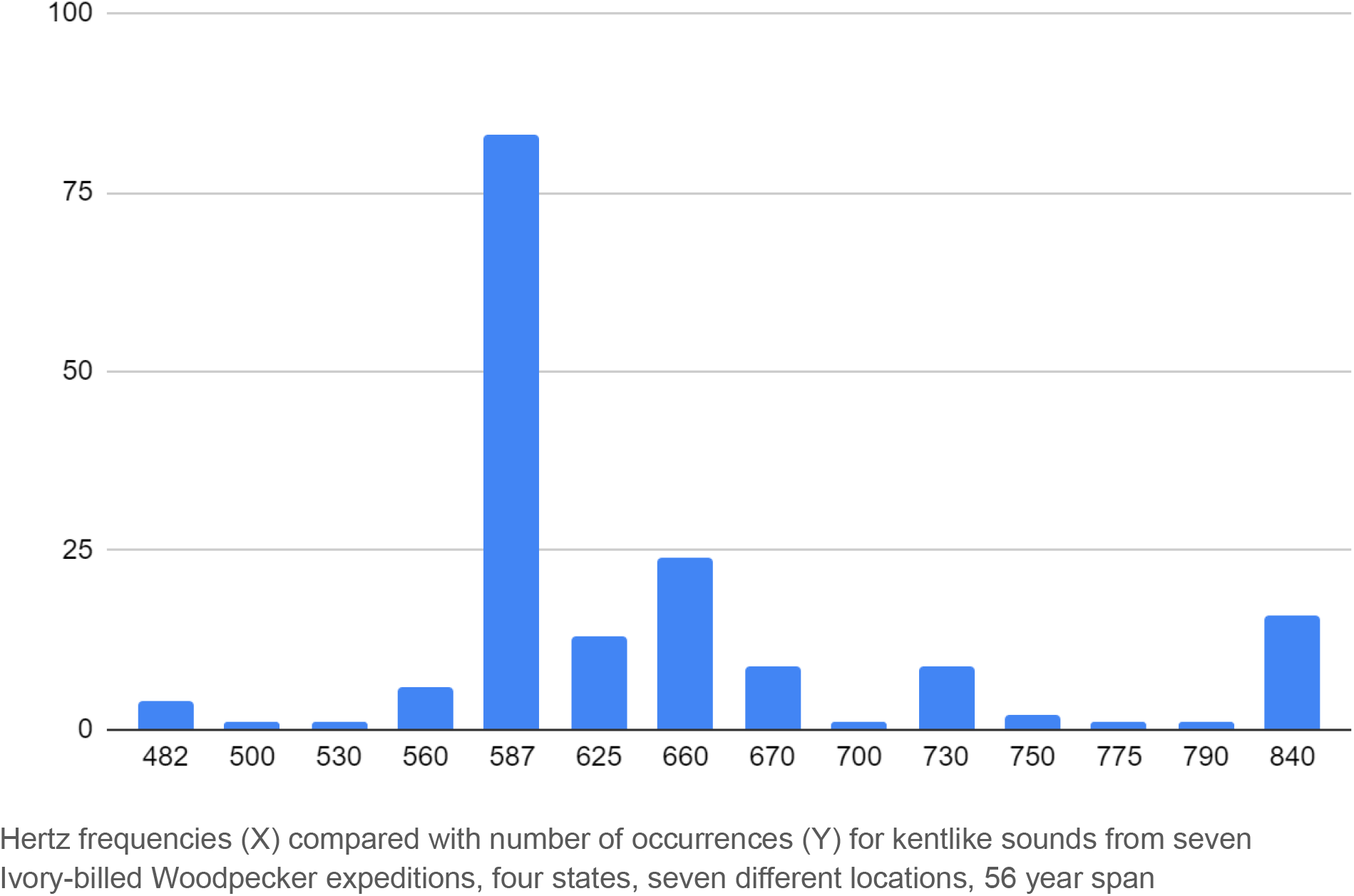

The results show a significant tendency for these sounds to be at 587 Hz (+/- 10 Hz), with a smaller spike, from the Cornell Arkansas recordings, around 660 Hz. There was one instance from the Cornell Arkansas recordings where a bird made a kent sound that slurred from 660 Hz down to 587 Hz. The spike at 840 Hz, largely recorded by Auburn-Windsor, may be Bluejay, although there are some indications that the IBWO can make kentlike sounds at and above this frequency [13] [19].

## Conclusion

The Ivory-billed Woodpecker had its population numbers reduced, perhaps drastically, in the late 1800s to early 1900s, through both hunting pressures and environmental degradation [1]. However, reports of its continued existence, including from focused scientific study, can be found in every decade since [1]. The idea that it is now extinct is perhaps a belief more of the uninformed. A student who becomes a specialist in this species will understand that the data and observations, both visual and acoustic, presented during this long period cannot be discounted.

Since the year 2000, with a number of major search efforts that were university-led in some cases, backed by more funding and better equipment, and with the advancement of technology for this equipment, more data has been obtained related to the Ivory-billed Woodpecker than at any time previous (save for the 1935 Allen-Kellogg Cornell expedition which was led to an active nest [2]). This of course would not be true for an extinct species.

It does not stand to reason that none of the recent data is valid from the premise that the species is extinct, and not reasonable to state that every single piece of the data, in many cases from professional ornithologists, is in error. It is more logical to presume that the growing level of technology is allowing us to get closer, so to speak, to a highly elusive, wary, and rare species. The data has value and should be examined objectively and quantitatively.

The idea of using spectrogram analysis to show fine detail of sounds is a more precise level of analysis than simply using the human ear, even an experienced one, as in earbirding. Spectrograms can verify what a human who has a “good musical ear” can say—there is harmony or dissonance, a sound is staccato or legato, it is a certain “note”, and even a particular individual. Importantly, spectrograms are used by government agencies for identification [20].

Discovering that Ivory-billed Woodpeckers make kent sounds near a specific Hz frequency makes sense. Virtually all organisms that make sounds have morphological constraints that are the cause. A working hypothesis is that the 587 Hz kent is produced by unstressed birds, unaware that they are being recorded. The 660 Hz kents recorded by Cornell in Arkansas may be an individual difference or a behavioral difference as in searching for a mate (speculative)– some reports from this expedition were of a lone bird [16].

Science is required to use logic. With this in mind–

1. If the sounds examined here are shown to be measurably different from other species, they are more likely IBWO.
2. If the sounds examined here compare well with other Campephilus, they are more likely IBWO.
3. If the sounds examined here are correlated with visual reports and images-videos from professional ornithologists, they are more likely IBWO.
4. If the sounds examined here, from different professional efforts, different years and locations, compare well with each other and reveal patterns, they are more likely IBWO.
5. If more than one of the above statements are true, the sounds are more likely IBWO.

All of the above logic statements are measurably true.

It is important to state here that, although science critique can demand “extraordinary claims require extraordinary evidence”, the level of evidence presented in this paper is quantitative and at a level higher than acceptable for identifying species by earbirding. Spectrograms describe more detail than earbirding can. Science cannot work with double standards, and perhaps with this in mind, the phrase “extraordinary claims require extraordinary evidence” might be amended to “extraordinary claims require a level of evidence sufficient to prove ordinary claims”. This is more pragmatic and workable. The idea of extraordinary evidence is also not a moving bar so to speak. Advances in technology and its precision allow what we might have in the past called extraordinary.

The author states that, for a non-biased and objective reader, the data analysis presented here is sufficient to show that the species, the rare Ivory-billed Woodpecker, continues to exist. Independent examination and expansion of the data presented is welcomed. Future data obtained can be compared with the results shown here and it is predicted that the Ivory-billed Woodpecker will be seen to use 587 Hz as a central point of its language.

## References

1. Haney JC. Woody’s last laugh: how the extinct Ivory-billed Woodpecker fools us into making 53 thinking errors. John Hunt Publishing (2021)

2. Allen, Arthur A and P. Paul Kellogg. Recent observations on the Ivory-billed Woodpecker. The Auk (1937)

3. Luneau, Guy. The Ivory-billed Woodpecker: taunting extinction: survival in the modern era. Zombie Media (2021)

4. Courtman, Matt. Personal communication.

5. Macaulay Library. Cornell Lab of Ornithology. https://www.macaulaylibrary.org/

6. Sounds: Is that an Ivory-billed Woodpecker? National Aviary. https://web.archive.org/web/20211018120417/ https://www.aviary.org/conservation/project-principalis/natural-history/sounds/

7. Sonic Visualizer. https://www.sonicvisualiser.org/

8. Xeno-canto. https://xeno-canto.org/

9. Online Tone Generator. https://www.szynalski.com/tone-generator/

10. Mennill, DJ. The search for the Ivory-billed Woodpecker: sounds. University of Windsor, Ontario, Canada. (2007)

11. Hill GE, Mennill DJ, Rolek B, Hicks TL, Swiston KA. Evidence suggesting that Ivory-billed Woodpeckers (Campephilus principalis) exist in Florida. Avian Conservation and Ecology. 1(3) (2006)

12. John Dennis. Ivory-billed Woodpecker recording (1968). Macaulay Library https://search.macaulaylibrary.org/catalog?taxonCode=ivbwoo

13. Collins, Michael D. Putative audio recordings of the Ivory-billed Woodpecker (Campephilus principalis). Acoustical society of America (2010)

14. Michaels, Mark A. 2023 in retrospect (part 4): acoustic data in a visual world. Project Principalis (formerly Project Coyote) Ivory-billed Woodpecker searches in Louisiana. Personal blog.

15. Chiandett, Cinzia and Giorgio Vallortigara. Chicks like consonant music. Psychological Science. October 22:10 (2011)

16. Charif, Russ. Ivory-billed Woodpecker Pages. Cornell University. Box. https://cornell.app.box.com/s/479gjn6lut74i4d4czwydilapomb0zd2

17. Personal communication and unpublished data.

18. Personal communication and unpublished data.

19. Webster, Rachel. Personal communication and unpublished data.

20. Spectrograph Mode for Voice ID. Baker Group—International (2023) https://www.bakerdvsa.com/spectrograph-mode-for-voice-id/

